# Sphingolipid metabolism-related genes as key regulatory hubs in white-smoke inhalation induced lung injury

**DOI:** 10.64898/2026.08.26.747407

**Authors:** Meng Fengji, Xin Haiming, Zhu xiulian, Cui Pei, Zhan Qiu, Li Rongsheng

## Abstract

**Objective:** White smoke inhalation injury (WSI) causes severe acute lung damage with no specific therapy currently available. Sphingolipid metabolism is implicated in pulmonary inflammation, but its transcriptional regulatory landscape in WSI remains unexplored. This study aimed to identify key sphingolipid metabolism-related genes and evaluate their regulatory roles and therapeutic potential in WSI.

**Methods:** We established a rat model of WSI and performed integrated bulk RNA-sequencing, weighted gene co-expression network analysis (WGCNA), and single-cell RNA sequencing (scRNA-seq) to screen for differentially expressed sphingolipid metabolism-related genes (DE-SRGs). Protein-protein interaction (PPI) network with four centrality algorithms was used to prioritize hub genes. In silico gene knockout and molecular docking were conducted to assess regulatory functions and identify potential drug candidates.

**Results:** We identified 22 DE-SRGs that were predominantly enriched in DNA replication and cell cycle pathways rather than canonical sphingolipid metabolic processes. PPI consensus prioritized three hub genes—*Top2a*, *Ttk*, and *Ccna2*—with *Top2a* exhibiting the highest expression in epithelial cells and significant downregulation after smoke exposure. ScRNA-seq revealed immune cell infiltration and epithelial differentiation trajectories. Virtual knockout showed that *Top2a* depletion affected the largest transcriptomic fraction (∼0.4%) and was enriched in lysosome biogenesis, innate immunity, phagocytosis, and lipid catabolism. Molecular docking identified thalidomide as a high-affinity ligand for *Top2a* (Vina score: −8.5 kcal/mol).

**Conclusion:** Our multi-omics integrative framework identifies *Top2a* as a central regulatory hub linking sphingolipid-associated inflammation to epithelial responses in WSI, and nominates thalidomide as a potential drug-repurposing candidate. These findings provide prioritized targets for future translational investigation.

## Introduction

Smoke bombs, which emit dense white smoke upon ignition, are widely used in fire drills and tactical evacuation training. Unlike smoke from wood or tobacco combustion, white smoke aerosols contain a complex mixture of corrosive toxic gases and particulate matter that can induce severe pulmonary injury upon inhalation[1,2]. Unintentional exposure to high concentrations in confined spaces may lead to irreversible lung fibrosis, acute respiratory distress syndrome (ARDS), and even mortality[3]. Toxicological studies have demonstrated that mixed toxicants in smoke bomb-derived aerosols collectively drive alveolar epithelial damage, inflammatory infiltration, and barrier disruption[4,5]. Currently, no specific therapeutics exist for white smoke inhalation-induced lung injury (WSI); clinical management is limited to symptomatic supportive care[6]. Therefore, elucidating the molecular pathological cascade of WSI and identifying actionable therapeutic targets remain urgent unmet needs.

Emerging evidence suggests that dysregulated lipid metabolism, particularly sphingolipid signaling, plays a pivotal role in the pathogenesis of acute lung injury. Sphingolipids are bioactive mediators that regulate epithelial survival, immune homeostasis, and inflammatory responses[7]. In toxic lung injury, excessive ceramide accumulation and disrupted sphingolipid metabolism are closely associated with chronic inflammation and loss of endothelial-epithelial barrier integrity[8,9]. For instance, sphingosine-1-phosphate receptor 1 (S1PR1) has been identified as a critical regulator in lipopolysaccharide(LPS) induced acute lung injury, where its downregulation exacerbates oxidative stress and pro-inflammatory cascades[10].However, the transcriptional landscape linking sphingolipid metabolic dysfunction to epithelial cellular responses specifically in WSI remains largely unexplored. More critically, existing transcriptomic studies on inhalation injury have been almost exclusively conducted using cigarette smoke, wood dust, or LPS models, while dedicated datasets for white smoke exposure are exceedingly scarce. Given the distinct chemical composition and pathogenic mechanisms of white smoke compared to other inhalation injury models, extrapolating findings across models may be misleading, severely hindering the elucidation of WSI-specific molecular pathology. Furthermore, previous studies have predominantly relied on bulk transcriptomics; the lack of single-cell resolution data further limits the precise characterization of transcriptional responses across distinct pulmonary cell populations in WSI.

To fill this research gap, we established rat models of white smoke inhalation and performed integrated bulk and single-cell transcriptomic profiling to systematically screen for genes co-altered with sphingolipid dysfunction. Specifically, we applied differential expression analysis, weighted gene co-expression network analysis (WGCNA), and protein-protein interaction (PPI) network topology to prioritize candidate hub genes. In parallel, we employed single-cell RNA sequencing to resolve cell-type-specific expression patterns, intercellular communication networks, and pseudotime differentiation trajectories. Furthermore, we conducted in silico gene knockout simulations and molecular docking to evaluate the regulatory potential of the identified hubs and to screen for existing compounds with therapeutic promise. By integrating these multi-level computational approaches, this study aims to establish a molecular framework that links sphingolipid metabolism to WSI pathology and to nominate viable targets for future translational intervention

## Materials and methods

### Animals and WSI model establishment

The Henan Skobes Biotechnology Co., LTD was the source of all the rats. SYXK(Gui)2023-0007 was the animal usage permission.All animal experimental procedures were approved by the 924th Hospital of the Joint Logistics Support Force of the Chinese PLA’s ethics commission (approval No. 924DWSY202405), and all animal handling and euthanasia operations strictly complied with standard animal ethical guidelines. Rats were kept in a room with constant ventilation, 12 hours of light and 12 hours of darkness, and a temperature of 25.0±1.0 °C with a relative humidity of 50.0±10.0%. They were also given free access to food and water.Referring to the rat model of white smoke inhalation (WSI) established by Cui et al.[11], the combustible materials in a small smoke-generating canister were placed in a multifunctional smoke injury experimental apparatus and ignited. After dense smoke had been generated, rats were immediately placed into a self-made highly permeable cage and exposed to the smoke environment for 5 min. The success of model establishment was verified by histopathological examination of lung tissues. The animal experiment included a normal group and a model group; rats were randomly assigned using a random number table into a control group (n = 8) and a model group (n = 10). The control group was placed in the self-made smoke experimental apparatus filled with air and allowed to breathe freely for 5 min, whereas the model group was subjected to the WSI model according to the above-described method.

### High-throughput RNA sequencing

High-throughput RNA sequencing of rat lung tissues was performed in collaboration with NovelBio Co., Ltd. Briefly, lung tissues were collected from rats, and cell viability and count were assessed during quality control. Total RNA was extracted, and its quality and quantity were evaluated. mRNA enrichment was conducted, followed by cDNA library construction with adapter and barcode incorporation for multiplexing. Library quality was verified, and sequencing was carried out on an Illumina platform. Raw reads were processed through adapter trimming, quality filtering, and alignment to a reference genome using tools such as STAR or HISAT, with gene expression levels quantified using HTSeq or featureCounts.

### Screening of Differentially Expressed SRGs (DE-SRGs) in WSI

After normalization, differential expression analysis between white smoke inhalation (WSI) and normal samples was performed using the limma R package. Differentially expressed genes (DEGs) were selected with thresholds of P < 0.05 and |log2FC| > 2, and visualized via volcano plots and heatmaps generated with the ggplot2 R package. Sphingolipid metabolism-related genes (SRGs) were retrieved from the GSEA-MSigDB database (https://www.gsea-msigdb.org/gsea/msigdb). Using the WGCNA R package, characteristic genes in core modules were extracted and intersected with DEGs and SRGs. The overlapping genes were considered candidate genes (designated as DE-SRGs) for subsequent analyses. Functional enrichment analyses were carried out using Gene Ontology (GO) and Kyoto Encyclopedia of Genes and Genomes (KEGG) databases.

### PPI Network Construction and Hub Gene Selection

A protein-protein interaction (PPI) network was constructed using the STRING database (https://cn.string-db.org/) to screen for hub genes among the target genes. The search was performed using the “Multiple Proteins by Names/Identifiers” mode, with the organism restricted to Rattus norvegicus and the minimum required interaction score set to 0.4. The full STRING interaction data were exported and visualized using Cytoscape software [43]. Subsequently, the cytoHubba plugin was employed to rank all network nodes based on four topological algorithms: Degree, EPC, MCC, and MNC. For each algorithm, genes ranked within the top 3 were retained as high-confidence candidates. To obtain robust consensus hub genes without imposing an overly stringent intersection, we applied a majority-voting strategy: Genes ranked top 3 by no less than three of the four topological algorithms were defined as consensus hub genes via majority-vote screening.

Drug candidates associated with the hub genes were retrieved from the Drug Signatures Database (DSigDB, https://dsigdb.tanlab.org/DSigDBv1.0/). To evaluate the binding affinity between the proteins encoded by the hub genes and the candidate drugs, molecular docking was performed. Specifically, the 3D chemical structures of non-toxic drugs with the highest interaction scores were obtained from the PubChem database (https://pubchem.ncbi.nlm.nih.gov/), while the spatial structures of the target proteins were acquired from AlphaFold (https://alphafold.ebi.ac.uk/). Docking simulations were subsequently carried out using the CB-Dock web server (https://cadd.labshare.cn/cb-dock/php/blinddock.php).

### Single-cell RNA sequencing

Single-cell RNA-seq experiments were performed by personnel at NovelBio Co., Ltd. Dissected tissues were preserved in dedicated storage solution, rinsed with RPMI-1640 medium, and minced into ∼1 mm³ fragments on ice. Tissue dissociation was achieved via enzymatic digestion with Collagenase I, Collagenase IV, and DNase I at 37 °C with continuous shaking for 40 min using a tissue dissociation device. The digested suspension was filtered through a cell strainer and centrifuged; cell pellets were resuspended in red blood cell lysis buffer to remove erythrocytes and washed repeatedly. Cell viability was evaluated by AO/PI staining using a fluorescence cell counter. Single-cell capture was performed on the NovelCyto Single-Cell Analysis System: cell suspensions were loaded into over 100,000 microwells containing barcoded oligonucleotide beads to achieve one bead per cell.

Following cell lysis, mRNA hybridized to bead-linked capture oligonucleotides; beads were then pooled for reverse transcription and Exo I digestion. Each cDNA fragment was labeled with unique molecular identifiers (UMIs) and cell barcodes to trace its cellular origin. Libraries were constructed following the standard whole-transcriptome amplification workflow, quantified, quality-checked, and subjected to paired-end 150 bp sequencing on a DNBSEQ-T7 platform.

Raw sequencing reads underwent adapter trimming, quality filtering, and genome alignment, with reads assigned to individual cells according to their barcodes. Post-sequencing quality control was carried out to exclude doublets and contaminants. Dimensionality reduction (PCA and t-SNE), unsupervised clustering, and differential expression analysis were performed using the Seurat R package to dissect cellular heterogeneity and identify cell-type marker genes. Cell trajectory and branch-gene analyses were implemented using the Monocle2 R package with DDRTree and default parameters, based on filtered expression matrices and cluster markers. Intercellular ligand-receptor interaction networks were inferred using the CellChat R package following Seurat normalization. In silico knockout simulation of hub genes was performed via the scTenifoldKnk [27]; Differentially expressed genes induced by gene perturbation were subjected to KEGG pathway enrichment analysis to identify significantly affected functional pathways.

### Statistical Analysis

Statistical analyses were conducted using R (version 4.3.0) and RStudio (version 2024.04.0+735). Unless otherwise indicated, statistical significance was defined as P < 0.05, with significance levels categorized as follows: *P < 0.05, **P < 0.01, ***P < 0.001, and ****P < 0.0001.

## Results

### Pathological changes in rat lung tissue after inhalation injury induced by white smoke

Hematoxylin-eosin (H&E) staining was performed to observe pulmonary pathological alterations in WSI rats (Fig 1). Lung tissue structure of the control group was clear and complete. At 72h post-WSI, Structural changes in rat lung tissue, infiltration of inflammatory cells, and congestion and bleeding.

**Fig 1.**
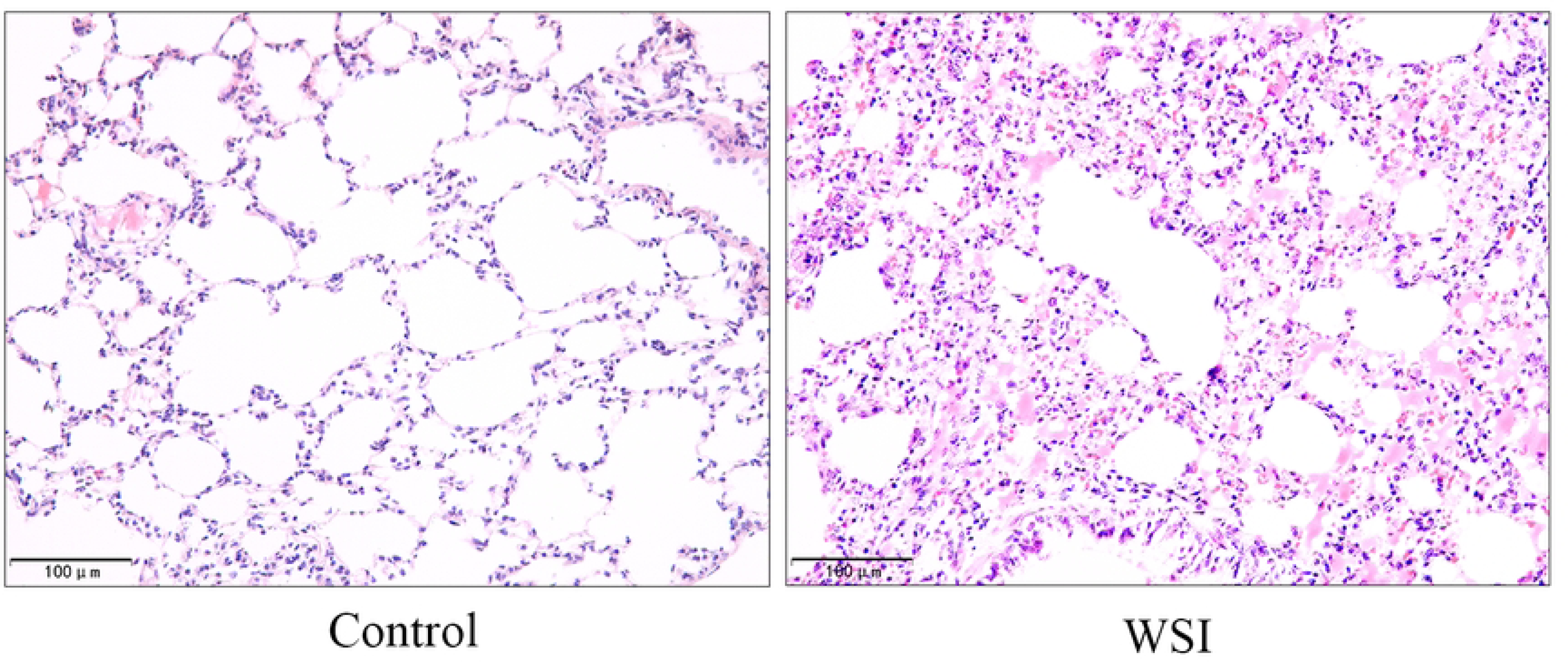
Histopathological changes of rat lung tissues following white smoke inhalation injury. **(A)** Representative images of H&E-stained lung sections from each group (Scale bar: 100 µm).

### Identification of DE-SRGs in WSI Vs Normal Samples

Sphingolipid metabolism-related genes(SRGs) were retrieved from the GSEA MSigDB (S1 Table). Differential expression analysis using limma package identified 971 differentially expressed genes (DEGs) between WSI and normal samples, comprising 620 upregulated and 351 downregulated genes (Fig. 2A and B). Weighted Gene Co-expression Network Analysis (WGCNA) was employed to identify core modules associated with WSI. A soft threshold of 24 was determined to be optimal, as it satisfied the scale-free topology criterion and ensured smooth connectivity (Fig. 2C). A cluster tree diagram of sample clustering is presented in Fig. 2D. Six distinct co-expression modules were identified (Fig. 2E), among which the blue module showed the strongest negative correlation with Cluster A (r = −0.98, P = 2 × 10^−12^) and the strongest positive correlation with Cluster B (r = 0.98, P = 2 × 10^−12^), yielding 1180 hub genes associated with WSI. To explore the role of sphingolipid metabolism in WSI pathology, we integrated DEGs, WGCNA hub genes, and SRGs via Venn diagram analysis, ultimately identifying 22 DE-SRGs for further investigation (Fig. 2F). Functional enrichment analysis revealed that these DE-SRGs were predominantly involved in DNA replication, nuclear chromosome segregation, and chromosome segregation (GO Biological Process; Fig. 2G). KEGG pathway analysis further indicated significant enrichment in Cell Cycle and Homologous Recombination pathways (Fig. 2H).

**Fig 2.**
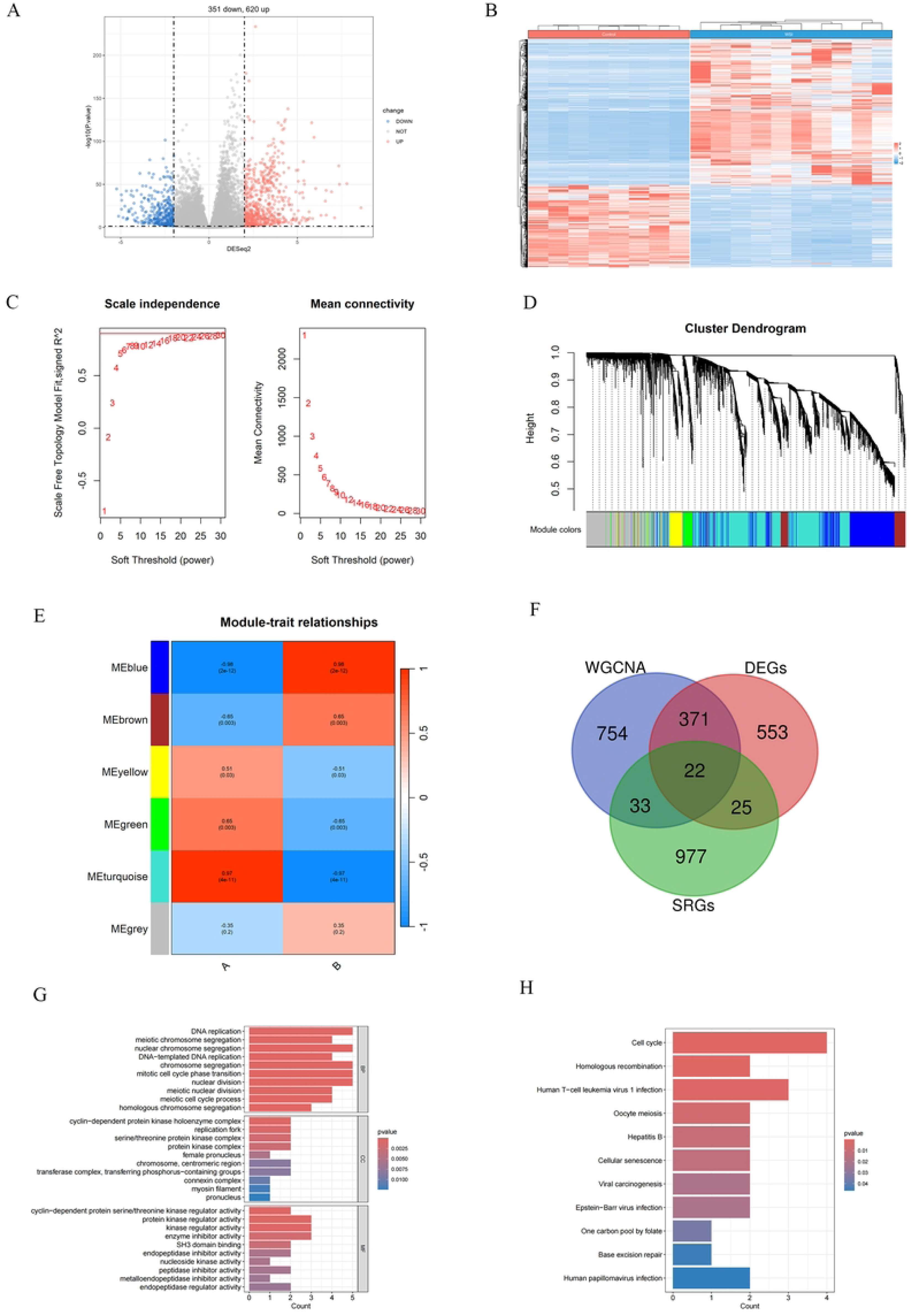
Screening and functional enrichment of DE-SRGs in WSI. **(A)** Volcano plot of all DEGs between the control and WSI groups. **(B)** Heatmap of all DEGs between the control and WSI groups (significance thresholds: p < 0.05 and |log₂FC| > 2). **(C)** Scale-free topology analysis for WGCNA to determine the optimal soft-thresholding power (β = 24). **(D)** Sample clustering dendrogram based on global gene expression profiles. **(E)** Heatmap displaying the correlation between six WGCNA co-expression modules and Cluster A/B scores; the blue module exhibited the strongest correlation with WSI-related traits. **(F)** Venn diagram showing the intersection of DEGs, WGCNA core module genes, and SRGs to identify DE-SRGs. **(G)** GO enrichment analysis of DE-SRGs in WSI. **(H)** KEGG pathway enrichment analysis of DE-SRGs in WSI.

### PPI network and hub gene analysis

To identify key disease-associated genes, DE-SRGs were mapped onto the STRING database to construct a protein-protein interaction (PPI) network, excluding unconnected nodes (Fig. 3A). Four centrality algorithms (Degree, EPC, MCC, and MNC) were applied to rank all nodes, retaining the top three candidates from each method. Through a majority voting consensus (genes appearing in the top 3 of at least three algorithms), we identified three hub genes: *Top2a*, *Ttk*, and *Ccna2*. As shown in the ranking heatmap (Fig 3B), these three genes consistently demonstrated high priority across multiple algorithms. In contrast, other candidates (*Tk1*, *Prr11*, *Arhgap11a*) were prioritized by only one or two algorithms, indicating less cross method consensus.

**Fig 3.**
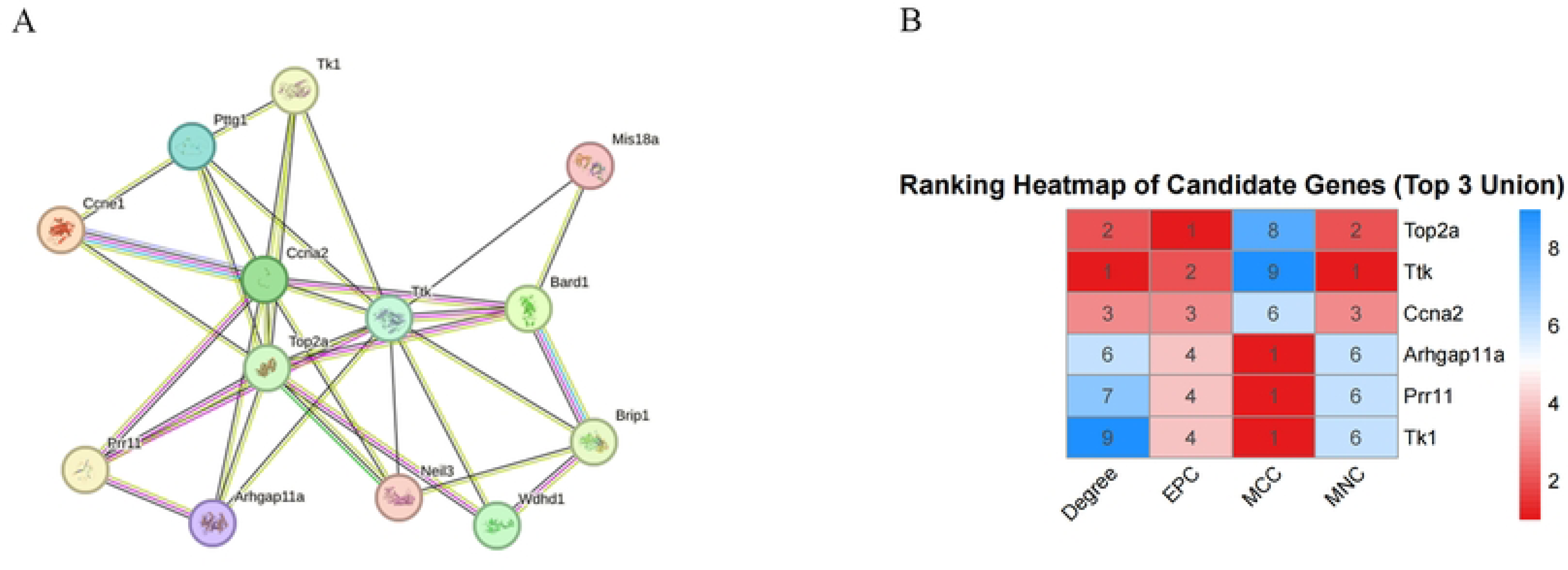
Consensus hub gene identification based on four PPI network topological algorithms. **(A)** PPI network of DE-SRGs.**(B)** Heatmap showing node rankings calculated by Degree, EPC, MCC and MNC algorithms.

### Single-Cell Transcriptomic analysis reveals cell-specific expression of the hub genes

Following the standard Seurat workflow, all cells were classified into 20 clusters based on the integrated RNA-seq and scRNA-seq datasets (Fig 4A). Cell annotation subsequently identified seven major cell types: T cells, B cells, NK cells, neutrophils, fibroblasts, macrophages, and epithelial cells (Fig 4B). In the Smoke group, the distribution density of T cells, B cells, NK cells, neutrophils, fibroblasts, and macrophages was markedly higher than that in the control group, whereas epithelial cells showed the opposite trend (Fig 4C), suggesting that the abundance of these immune and stromal cell types increases upon smoke exposure.

**Fig 4.**
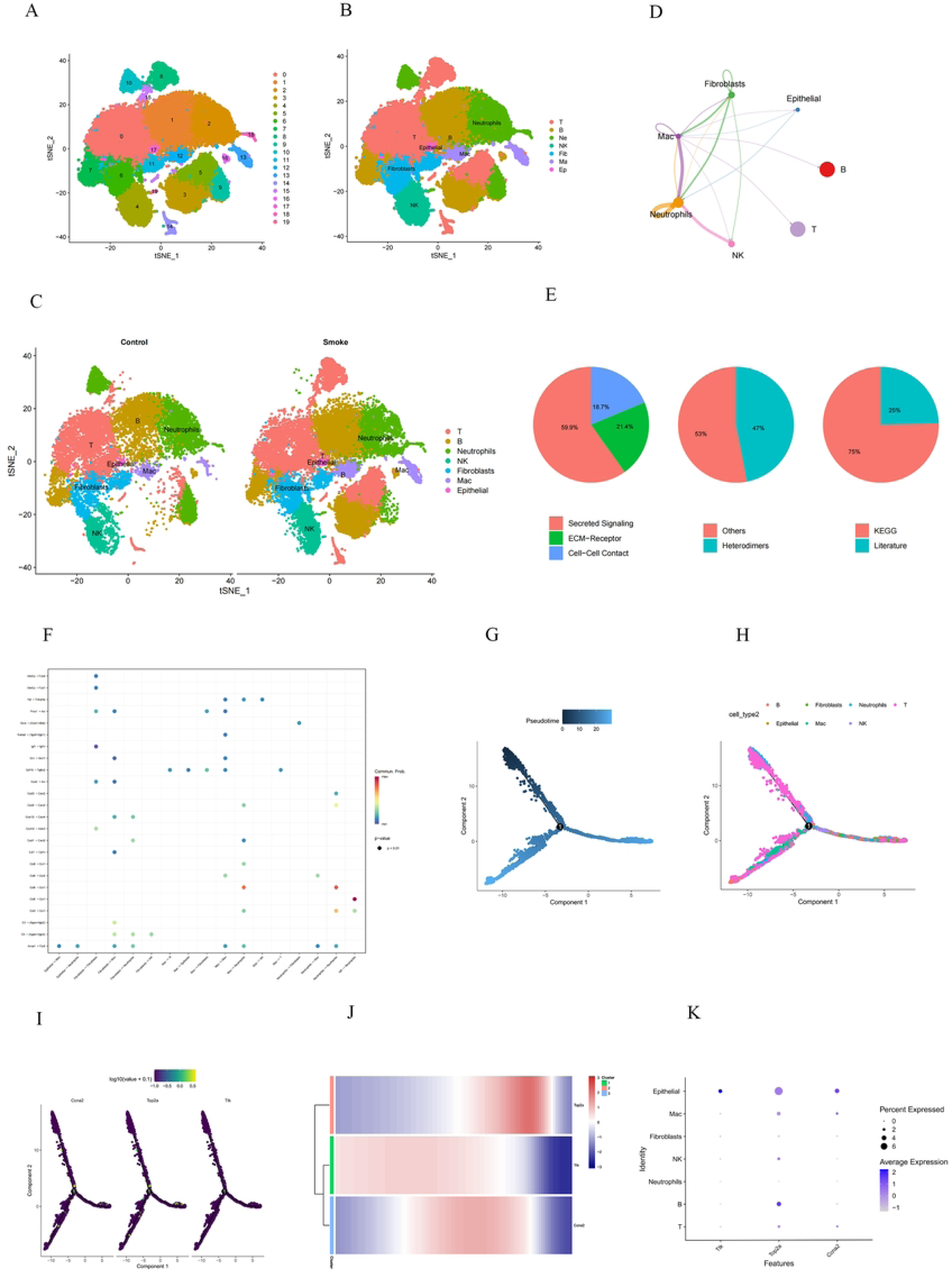
Single-cell transcriptomic characterization of WSI lung tissue. **(A)** t-SNE plot of cellular clusters. **(B)** t-SNE plot colored by cell type. **(C)** Split t-SNE plots grouped by sample condition. **(D)**Pie chart showing the proportion of intercellular signaling modes, including secreted signaling, ECM-receptor interaction, and cell-cell contact, based on CellChat analysis. **(E)**Network diagram of differential intercellular communication intensity (red: enhanced; blue: attenuated).**(F)** Dot plot showing significant ligand–receptor interactions in receiver cells. Dot size and color indicate communication strength. **(G)** Pseudotime trajectory colored by differentiation stage (dark = early; light = late). **(H)** Pseudotime trajectory colored by cell type. **(I)** Feature plots showing pseudotime-dependent expression dynamics of key genes. **(J)**Heatmap depicting dynamic expression trends of key genes along pseudotime. **(K)** Bubble plot displaying average expression level and expression percentage of key genes across all cell types.

CellChat analysis was performed to evaluate intercellular communication within the pulmonary microenvironment. Signaling networks comprised secreted signaling (59.9%), cell-cell contacts (21.4%), and ECM-receptor interactions (18.7%) (Fig 4D). Ligand-receptor pair characterization revealed that 53% were mediated by monomeric/non-canonical complexes and 47% by heterodimeric complexes. Approximately 75% of signaling pathways were derived from the KEGG database, with the remainder curated from literature. Myeloid cells (macrophages and neutrophils) and fibroblasts emerged as central communication hubs(Fig 4E). Specifically, Fibroblasts targeted macrophages and Neutrophils via C3-ITGAM/ITGB2 and Cxcl12-Cxcr4 axes, and macrophages specifically via Gas6/Pros1-Axl pathways. Conversely, macrophages regulated Fibroblasts and Epithelial cells through the Gdf15-Tgfbr2 axis and counter-regulated neutrophils via Ccl/Cxcl family members and the Tnf-Tnfrsf1b pathway, establishing bidirectional crosstalk between immune and stromal compartments(Fig 4F).

Pseudotime trajectory analysis, with cells colored by differentiation stage (dark = early; light = late), revealed a bifurcated cellular response to smoke exposure (Fig. 4G). The left branch was enriched with T cells, macrophages, and neutrophils, representing an immune-inflammatory activation axis. The right branch was dominated by fibroblasts, indicating a tissue remodeling and fibrotic progression axis. B cells, NK cells, and epithelial cells were distributed around the bifurcation point, suggesting their involvement in fate determination(Fig. 4H). Trajectory-dependent expression analysis showed that *Top2a* and *Ccna2* were significantly upregulated at the bifurcation point and the terminal end of the right branch (Fig. 4I), coinciding with fibroblast enrichment. *Top2a* peaked at intermediate-to-late pseudotime stages before gradually declining (Fig. 4J). Collectively, these findings suggest that smoke inhalation induces a biphasic pulmonary response: an early immune-inflammatory phase followed by a proliferative/fibrotic phase characterized by upregulated cell cycle machinery in fibroblasts. At the single-cell level, *Top2a* exhibited the highest average expression and largest positive fraction in epithelial cells, with moderate expression in macrophages, NK cells, B cells, and T cells (Fig. 4K).

### Transcriptomic Alterations Following Virtual hub genes Knockout

To evaluate the transcriptional regulatory capacities of *Ttk*, *Top2a*, and *Ccna2*, we performed in silico knockout using scTenifoldKnk and compared the perturbed transcriptomes. Depletion of *Top2a* shifted expression of approximately 0.4% of all genes, a larger fraction than that for *Ttk* (approximately 0.3%) and *Ccna2* (approximately 0.1%) (Fig 5A and B), indicating that *Top2a* exerts the greatest global transcriptional regulation among the three. KEGG enrichment of knockout of key genes further supported this (Fig 5C): those from *Top2a* knockout were prominently enriched in lysosome biogenesis, tuberculosis infection, innate immunity, lipid catabolism, and phagocytosis. *Ttk* perturbation showed largely overlapping pathway signatures, with additional enrichment in efferocytosis, whereas *Ccna2*-associated differentially expressed genes were exclusively linked to ribosome translation and amino acid metabolism, lacking any immune-related terms. Consistently, multi-layer GO enrichment (Biological Process, Cellular Component, Molecular Function) (Fig 5C) validated *Top2a*’s regulatory roles in immune inflammation, phagocytosis, lipid metabolism, and lysosome assembly, with corresponding subcellular (lysosomes, vesicles) and molecular (cytokines, lipid transporters, hydrolases) annotations. *Ttk* shared most GO terms with *Top2a*, while *Ccna2* was only enriched in basic protein synthesis-related processes.

**Fig 5.**
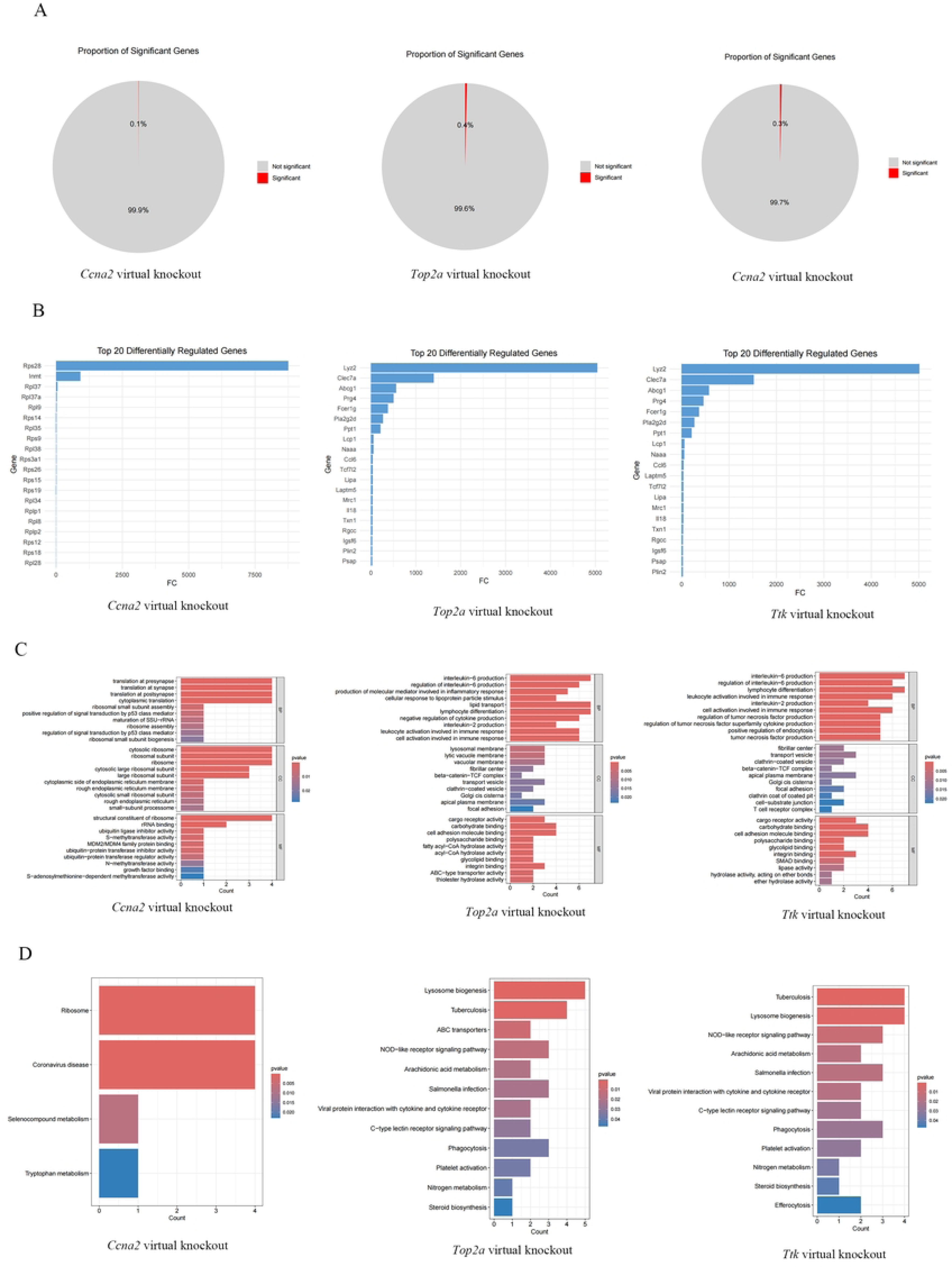
Transcriptional regulatory impact of key gene virtual knockout in WSI lung tissue. **(A)** Proportion of significantly differentially expressed genes in the whole transcriptome after key gene virtual knockout. **(B)** Barplot of top 20 differentially regulated genes knockout of key genes respectively **(C)** KEGG pathway enrichment analysis of differentially regulated genes following individual key gene perturbation. **(D)** Multi-layer GO enrichment analysis validating functional roles of differentially regulated genes associated with each key gene knockout.

### Molecular Docking Simulation of Candidate Drugs Targeting Key Genes

To screen candidate therapeutic agents targeting the hub gene *Top2a*, we first conducted drug enrichment analysis based on DSigDB, and nine compounds predicted to interact with all three hub genes were obtained (Figure 6A). We further filtered non-toxic candidates with high binding affinity scores and publicly available 3D structures from the PubChem database, among which thalidomide was selected for subsequent molecular docking validation. Docking results demonstrated that the Vina binding energies of thalidomide against all three hub proteins were lower than −6 kcal/mol (Table 1), representing stable intermolecular binding. Of note, *Top2a* protein displayed the highest binding affinity to thalidomide, with a Vina score of −8.5 kcal/mol. Collectively, thalidomide was determined as the optimal therapeutic candidate for WSI, and the simulated drug-protein binding conformations are displayed in Fig 6B.

**Fig 6.**
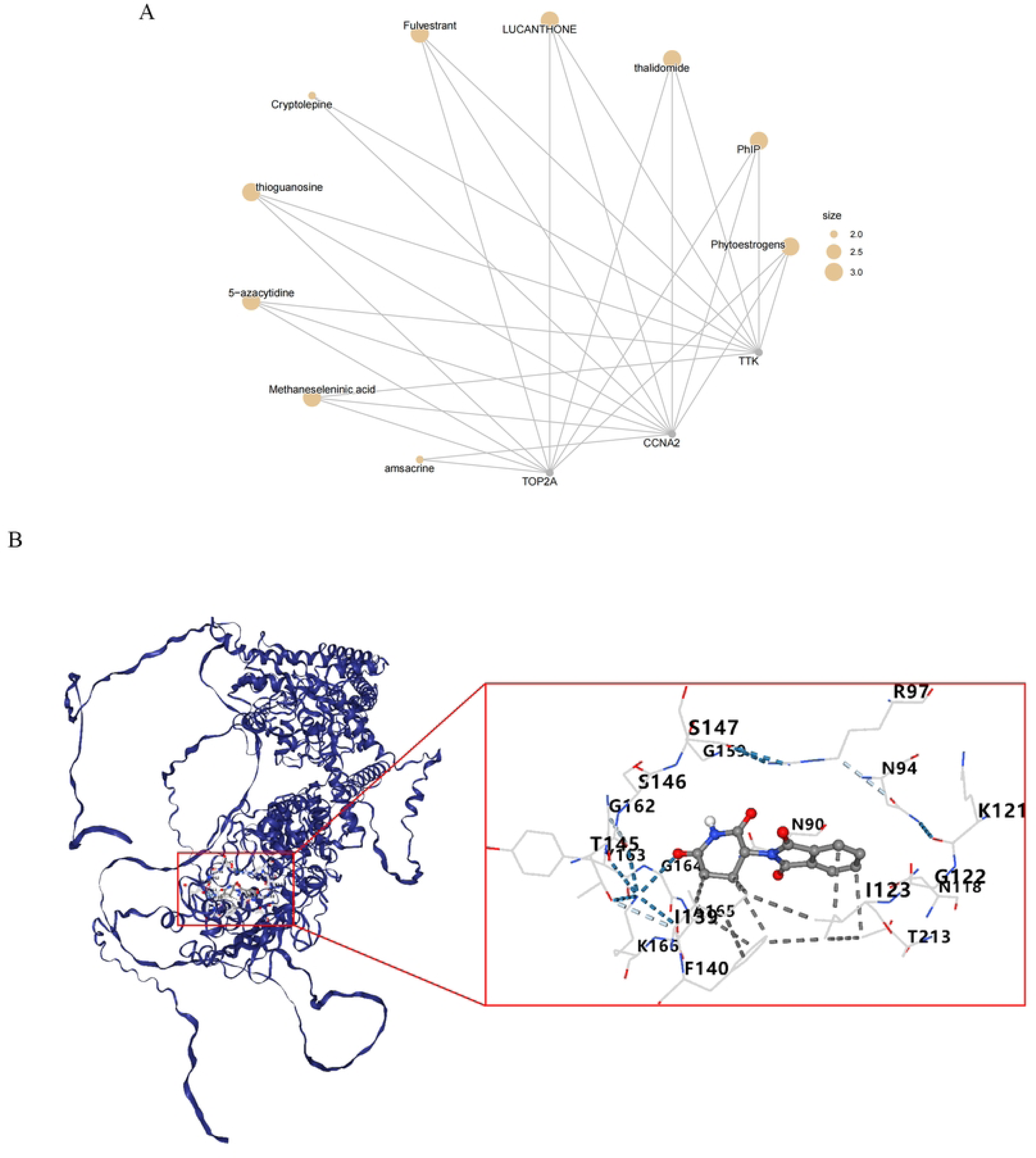
Drug enrichment analysis and molecular docking validation for WSI hub genes. **(A)** Cnet plot of the relationship between the most significant drug terms and their associated key genes. Nodes are arranged in a circular layout; drug-related terms are represented as larger nodes, and individual genes as smaller nodes. Edges connect genes to their corresponding drug annotations based on DSigDB enrichment analysis. **(B)** Docking of *Top2a* with thalidomide.

**Table 1.** The binding energy between key genes and drugs.

| Gene | AlphaFoldDB ID | DRUG | PubChem CID | Binding energy (kcal/mol) |
| --- | --- | --- | --- | --- |
| *Ccna2* | AF-A0JPK0-F1 | thalidomide | 5426 | -6.4 |
| *Ttk* | AF-A0A8I5ZNI6-F1 | thalidomide | 5426 | -7.9 |
| *Top2a* | AF-P41516-F1 | thalidomide | 5426 | -8.5 |

## DISCUSSION

In the present study, we systematically delineated the transcriptomic landscape of white smoke inhalation-induced lung injury (WSI) by integrating bulk RNA sequencing, weighted gene co-expression network analysis (WGCNA), single-cell transcriptomics, computational gene perturbation simulation, and molecular docking. Our findings not only corroborated the pathological hallmarks of WSI—including disruption of pulmonary tissue architecture, inflammatory cell infiltration, and congestion with hemorrhage—but also established, to our knowledge for the first time in this pathological context, a functional molecular link between sphingolipid metabolism dysregulation and specific transcriptional hubs.

Notably, the 22 differentially expressed sphingolipid metabolism-related genes (DE-SRGs) identified in this study were predominantly enriched in DNA replication, chromosome segregation, and cell cycle regulatory pathways (Figures 2F and 2G), rather than in canonical sphingolipid metabolic or inflammatory signaling cascades. This unexpected enrichment profile suggests that under WSI stress conditions, transcriptional dysregulation of sphingolipid metabolism-related genes primarily impairs epithelial cell proliferation and genomic stability, rather than directly mediating classical inflammatory signaling. This finding resonates with recent studies: Hannun and Obeid[12] proposed that sphingolipid metabolites (such as ceramide and sphingosine-1-phosphate) can influence the balance between proliferation and apoptosis through cell cycle checkpoint regulation; meanwhile, Jamjoum R et al. [13] demonstrated that alterations in sphingolipid metabolic enzyme expression can indirectly affect the transcriptional activity of DNA damage repair-related genes. Furthermore, Bi W et al. [14] found that aberrant activation of cell cycle-related genes constitutes a critical step in epithelial repair failure in acute lung injury models. Therefore, our results may reveal a previously underappreciated pathological mechanism: WSI-induced sphingolipid metabolic disturbance compromises the regenerative capacity of pulmonary epithelial cells by interfering with cell cycle progression, thereby exacerbating the irreversibility of lung injury. By identifying transcriptional candidates potentially involved in such metabolic derangements, our study extends prior observations and provides novel entry points for subsequent targeted interventions.

Single-cell RNA sequencing analysis further unveiled profound remodeling of pulmonary cellular composition following smoke exposure: immune cells (T cells, B cells, NK cells, neutrophils, and macrophages) and fibroblasts were significantly expanded, while epithelial cells were markedly diminished. This immune-stromal enrichment pattern is consistent with the acute inflammatory response characteristic of inhalation injury [15]. Mercel AI et al. [16] similarly observed early and massive recruitment of alveolar macrophages and neutrophils in a murine model of smoke inhalation injury, with the intensity of immune cell infiltration positively correlated with injury severity. Our data further indicate that the significant expansion of fibroblasts may herald the early initiation of fibrotic processes, a finding concordant with the fibroblast activation trajectory identified by Liu W et al. [17] in bleomycin-induced pulmonary fibrosis models.

Of particular importance, pseudotime trajectory analysis revealed a bifurcated cellular response upon smoke insult: the left branch represented an immune-inflammatory axis enriched in myeloid immune cells, whereas the right branch corresponded to a tissue-remodeling and fibrotic progression axis dominated by fibroblasts. Epithelial cells, together with B cells and NK cells, were localized predominantly around the bifurcation point, implying that these cell populations are situated at the critical fate-decision checkpoint following smoke-induced lung injury. Trajectory-dependent gene expression demonstrated that *Top2a* and *Ccna2* were markedly upregulated at the bifurcation point and at the terminal end of the fibroblast-enriched right branch. Collectively, these trajectory profiles support a biphasic pathological cascade after WSI: an early immune-inflammatory phase followed by a later proliferative and fibrotic remodeling phase accompanied by heightened cell-cycle-related transcriptional activity.

Although our pseudotime results do not provide direct evidence for epithelial-to-mesenchymal phenotypic conversion along trajectory branches, epithelial plasticity is widely recognized as a key driver that promotes both fibrotic remodeling and persistent inflammation in acute lung injury [17,18] constituting a “dual threat” in WSI pathology. EMT is well-established as one critical pathogenic module in pulmonary fibrosis [19]. Notably, single-cell profiling demonstrated that *Top2a* exhibited the highest average expression and positive fraction within epithelial cells, with moderate expression in multiple immune cell types. The elevated cell-cycle-related gene signature at the bifurcation node and fibrotic-branch terminus suggests that cell-cycle machinery activation may coordinate multi-cellular fate decisions across epithelial and stromal compartments during WSI-provoked lung repair and fibrosis. This observation provides a novel perspective for understanding the early molecular events underlying post-WSI pulmonary fibrosis.

Among the three hub genes—*Top2a*, *Ttk*, and *Ccna2*—prioritized via consensus screening from the PPI network, we observed distinct cell-type-dependent expression patterns and divergent functional characteristics. Consistent with our single-cell profiling (Fig.4K), *Top2a* displayed the highest expression level in epithelial cells, and its expression was significantly downregulated upon smoke exposure; *Ttk* showed an increasing expression trend after injury, whereas *Ccna2* exhibited a non-significant upward tendency. Such cell-specific expression signatures indicate that these three hub genes exert their biological functions within distinct pulmonary cellular compartments. More importantly, our computational gene-knockout simulations uncovered their disparate regulatory landscapes. Depletion of *Top2a* perturbed ∼0.4% of the overall transcriptome, with significant enrichment of pathways related to lysosome biogenesis, innate immunity, phagocytosis, and lipid catabolism. These biological processes are essential for alveolar macrophage function, pulmonary surfactant homeostasis, and epithelial host defense [20,21]. Dysfunction of the autophagy-lysosomal axis has been well-documented in multiple chronic pulmonary diseases, including chronic obstructive pulmonary disease and idiopathic pulmonary fibrosis [22]. Of particular note, perturbation of *Top2a* strongly altered lipid catabolic pathways. Lipid metabolic derangements, particularly accumulation of phospholipids and sphingolipids, are known to impair surfactant function and further aggravate lung injury [23]. These observations support *Top2a* as a central molecular node linking sphingolipid metabolic dysfunction to pathological progression in WSI. Perturbation of *Ttk* shared multiple immune- and lipid-associated pathways with *Top2a*, but additionally enriched processes involved in efferocytosis [24] Efferocytosis, the clearance of apoptotic cells, is indispensable for resolving inflammatory responses; impaired efferocytosis perpetuates inflammation and promotes fibrotic progression [25]. Therefore, *Ttk* may serve as an auxiliary modulator facilitating inflammation resolution following white-smoke-induced lung damage. In contrast, *Ccna2* knockout mainly disturbed ribosomal translation and amino-acid metabolism without enriching immune- or inflammation-related functional terms. This suggests that within this WSI pathological context, *Ccna2* likely acts primarily as a housekeeping gene rather than a major pathological driver. Collectively, this functional stratification positions *Top2a* as the dominant regulatory hub connecting sphingolipid-associated intracellular signaling to multicellular tissue-level responses, with *Ttk* playing secondary overlapping roles in immune regulation, and *Ccna2* exerting limited pathological contribution. These findings provide critical evidence for prioritizing candidate targets for future therapeutic intervention.

Cell-cell communication analysis further expands this mechanistic framework. Based on ligand-receptor interaction predictions, we identified macrophages and fibroblasts as core signaling hubs, while neutrophils represented the primary responsive target cells. Specifically, fibroblasts signal to macrophages and neutrophils via the C3-ITGAM/ITGB2 and Cxcl12-Cxcr4 axes, and regulate macrophages through the Gas6/Pros1-Axl pathway. Macrophages, in turn, act on fibroblasts and epithelial cells via the Gdf15-Tgfbr2 axis and reciprocally communicate with neutrophils through Ccl/Cxcl family chemokines and the Tnf-Tnfrsf1b pathway, forming bidirectional crosstalk between immune and stromal compartments. This complex intercellular signaling landscape resembles observations reported in bronchoalveolar lavage samples from patients with severe ARDS[26]. Critically, paracrine signals originating from fibroblasts and macrophages are predicted to upregulate *Top2a* expression in epithelial cells, which may amplify epithelial-derived inflammatory responses. This finding indicates that *Top2a* is not merely an intracellular effector molecule, but also a downstream integrator of microenvironmental cues. We thus propose a microenvironment-*Top2a*-epithelial response axis that bridges extracellular immune-stromal crosstalk and intracellular sphingolipid-dependent epithelial dysfunction. This axis offers new mechanistic insights into cascade amplification during WSI-mediated lung injury.

Given the pivotal regulatory status of *Top2a*, we explored its therapeutic potential through drug enrichment analysis and molecular docking. Among nine drugs predicted to interact with all three hub genes, we selected thalidomide—a clinically available immunomodulatory agent with well-established anti-inflammatory and anti-angiogenic properties [27,28]—for docking simulation. The Vina score of −8.5 kcal/mol between thalidomide and *Top2a* indicated superior binding affinity compared to its interactions with *Ttk* and *Ccna2*, suggesting that *Top2a* represents the primary target among the three. The anti-inflammatory mechanisms of thalidomide have been extensively investigated: it suppresses pro-inflammatory cytokine production (including TNF-α) through inhibition of the NF-κB signaling pathway[29], exerts anti-angiogenic effects by inhibiting vascular endothelial growth factor (VEGF) [30], and modulates adaptive immune responses by regulating T cell subset balance [31]. Notably, thalidomide is known to modulate macrophage polarization [32], and macrophage phenotypic switching is tightly governed by lipid-metabolism-associated transcriptional reprogramming [33]. This observation intriguingly echoes the enrichment of lipid catabolic pathways observed following virtual *Top2a* knockout in our study.Moreover, thalidomide has demonstrated certain anti-fibrotic effects in animal models of pulmonary fibrosis [34], providing additional theoretical support for its potential application in fibrotic processes that may ensue following WSI. However, the precise mechanism of interaction between thalidomide and *Top2a*—whether through enzymatic activity inhibition, transcriptional interference, or allosteric regulation—remains to be experimentally validated. Furthermore, the known adverse effects of thalidomide, including teratogenicity and neurotoxicity [35], warrant careful consideration in clinical translation; future efforts may need to develop its structural analogs (such as lenalidomide or pomalidomide) to optimize the efficacy-safety balance.

Several limitations should be acknowledged. First, all phenotypic and molecular conclusions of this study are obtained from rat WSI models and computational bioinformatics predictions. Further validation using human lung tissues or primary pulmonary epithelial cells is required to confirm its translational value for clinical human WSI injury. Second, the in silico knockout and docking results are hypothesis-generating rather than confirmatory; future studies employing CRISPR-Cas9-mediated gene editing, pharmacological inhibition, and functional rescue experiments are necessary to establish causality. Third, while we identified *Top2a* as a central hub, the interplay between *Top2a* and specific sphingolipid species (e.g., ceramide, S1P) was not directly measured and warrants targeted lipidomic analysis. Despite these limitations, our multi-omics integrative framework provides a robust foundation for understanding WSI pathogenesis and prioritizes actionable targets for subsequent investigation.

## Data availability

Bulk RNA-seq and scRNA-seq raw data of this study are available in the GEO database under accession numbers GSE341464 and GSE341719, respectively.

## Supporting information

S1 Table. Sphingolipid metabolism-related genes were retrieved from the GSEA MSigDB.

## References

1. Schaeffer DJ, Kapila S, Meadows JE, Hinderberger E, Wentsel R. Chemical characterization of residues from military HC smokepots. J Hazard Mater. 1988;17: 315–328. doi:10.1016/0304-3894(88)85007-6

2. Klapötke T, Shaw A, Glueck J. Effect of adding 5-aminotetrazole to a modified U.S. army terephthalic acid white smoke composition. Cent Eur J Energ Mater. 2017;14: 489–500. doi:10.22211/cejem/76843

3. Murray BP, Ralston SA, Dunkley CA, Carpenter JE, Geller RJ, Kazzi Z. Pneumonitis and respiratory failure secondary to civilian exposure to a smoke bomb in a partially enclosed space. J Spec Oper Med Peer Rev J SOF Med Prof. 2018;18: 24–26. doi:10.55460/UD9X-AUXA

4. Reczyńska K, Tharkar P, Kim SY, Wang Y, Pamuła E, Chan H-K, et al. Animal models of smoke inhalation injury and related acute and chronic lung diseases. Adv Drug Deliv Rev. 2018;123: 107–134. doi:10.1016/j.addr.2017.10.005

5. Antonio ACP, Castro PS, Freire LO. Smoke inhalation injury during enclosed-space fires: an update. J Bras Pneumol Publicacao Of Soc Bras Pneumol E Tisilogia. 2013;39: 373–381. doi:10.1590/S1806-37132013000300016

6. Raymond LW. Fire-related inhalation injury. N Engl J Med. 2016;375: 1904. doi:10.1056/NEJMc1611256

7. Ghidoni R, Caretti A, Signorelli P. Role of sphingolipids in the pathobiology of lung inflammation. Mediators Inflamm. 2015;2015: 487508. doi:10.1155/2015/487508

8. Petrache I, Berdyshev EV. Ceramide signaling and metabolism in pathophysiological states of the lung. Annu Rev Physiol. 2016;78: 463–480. doi:10.1146/annurev-physiol-021115-105221

9. Zhang Y, Zhang J, Fu Z. Molecular hydrogen is a potential protective agent in the management of acute lung injury. Mol Med. 2022;28: 27. doi:10.1186/s10020-022-00455-y

10. Rojas-Quintero J, Polverino F. Molecular maestro in the lung: the sphingolipid rheostat. Am J Respir Cell Mol Biol. 2025;73: 170–172. doi:10.1165/rcmb.2025-0067ED

11. Cui P, Xin H, Yao Y, Xiao S, Zhu F, Gong Z, et al. Human amnion-derived mesenchymal stem cells alleviate lung injury induced by white smoke inhalation in rats. Stem Cell Res Ther. 2018;9: 101. doi:10.1186/s13287-018-0856-7

12. Hannun YA, Obeid LM. Principles of bioactive lipid signalling: lessons from sphingolipids. Nat Rev Mol Cell Biol. 2008;9: 139–150. doi:10.1038/nrm2329

13. Jamjoum R, Majumder S, Issleny B, Stiban J. Mysterious sphingolipids: metabolic interrelationships at the center of pathophysiology. Front Physiol. 2023;14: 1229108. doi:10.3389/fphys.2023.1229108

14. Bi W, Wang R, Hou S, Liu W, Zhang H, Xu Z, et al. Alveolar epithelial type 2 cells in acute lung injury and acute respiratory distress syndrome: spatiotemporal fate and regulation. Burns Trauma. 2025;13: tkaf050. doi:10.1093/burnst/tkaf050

15. Mercel A, Tsihlis ND, Maile R, Kibbe MR. Emerging therapies for smoke inhalation injury: a review. J Transl Med. 2020;18: 141. doi:10.1186/s12967-020-02300-4

16. Mercel AI, Gillis DC, Sun K, Dandurand BR, Weiss JM, Tsihlis ND, et al. A comparative study of a preclinical survival model of smoke inhalation injury in mice and rats. Am J Physiol Lung Cell Mol Physiol. 2020;319: L471–L480. doi:10.1152/ajplung.00241.2020

17. Liu W, Meridew JA, Aravamudhan A, Ligresti G, Tschumperlin DJ, Tan Q. Targeted regulation of fibroblast state by CRISPR-mediated CEBPA expression. Respir Res. 2019;20: 281. doi:10.1186/s12931-019-1253-1

18. Planté-Bordeneuve T, Pilette C, Froidure A. The epithelial-immune crosstalk in pulmonary fibrosis. Front Immunol. 2021;12: 631235. doi:10.3389/fimmu.2021.631235

19. Han Y-Y, Shen P, Chang W-X. Involvement of epithelial-to-mesenchymal transition and associated transforming growth factor-β/smad signaling in paraquat-induced pulmonary fibrosis. Mol Med Rep. 2015;12: 7979–7984. doi:10.3892/mmr.2015.4454

20. Barna BP, Judson MA, Thomassen MJ. Inflammatory pathways in sarcoidosis. Adv Exp Med Biol. 2021;1304: 39–52. doi:10.1007/978-3-030-68748-9_3

21. McElroy MC, Kasper M. The use of alveolar epithelial type I cell-selective markers to investigate lung injury and repair. Eur Respir J. 2004;24: 664–673. doi:10.1183/09031936.04.00096003

22. Kim H, Wang W, Dobrescu I, Lee J, Martorelli J, Wang S, et al. Autophagy in the lung: guardian of homeostasis or driver of disease. Autophagy Rep. 2025;4: 2568537. doi:10.1080/27694127.2025.2568537

23. Xu K, Shao X, Lu R, Liao Y, Zhao Y, Wang B, et al. Effective-component compatibility of bufei yishen formula III protects lung air-blood barrier by regulating the oxidative stress: via the nuclear factor-E2-related factor 2 pathway. Int J Chron Obstruct Pulmon Dis. 2025;Volume 20: 2211–2226. doi:10.2147/COPD.S513071

24. Sadaf S, Zhang X, Lei L, Shen J, Jamal I, Modugu G, et al. Macrophage efferocytosis is controlled by epigenetic modifications mediated by RBPJ. Immunology; 2025. doi:10.1101/2025.10.10.681723

25. Kirtay M, Ispirjan M, Bonnard B, Bruggner A-L, Boehringer L, Miessler M, et al. CD47 blockade reprograms the monocyte-macrophage axis to promote inflammation resolution in atherosclerosis. bioRxiv: The Preprint Server for Biology; 2026. doi:10.64898/2026.04.24.720546

26. Asaba CN, Bitazar R, Labonté P, Bukong TN. Bronchoalveolar lavage single-cell transcriptomics reveals immune dysregulations driving COVID-19 severity. PloS One. 2025;20: e0309880. doi:10.1371/journal.pone.0309880

27. Franks ME, Macpherson GR, Figg WD. Thalidomide. The Lancet. 2004;363: 1802–1811. doi:10.1016/S0140-6736(04)16308-3

28. Nikovia K, Kapsalis M, Georgoulakis M, Panousis A, Neochoritis CG. The multifaceted legacy of thalidomide: chemistry and biology driving modern drug design. ChemMedChem. 2026;21: e202501105. doi:10.1002/cmdc.202501105

29. Moreira AL, Sampaio EP, Zmuidzinas A, Frindt P, Smith KA, Kaplan G. Thalidomide exerts its inhibitory action on tumor necrosis factor alpha by enhancing mRNA degradation. J Exp Med. 1993;177: 1675–1680. doi:10.1084/jem.177.6.1675

30. Karajeh MA, Hurlstone DP, Stephenson TJ, Ray-Chaudhuri D, Gleeson DC. Refractory bleeding from portal hypertensive gastropathy: a further novel role for thalidomide therapy? Eur J Gastroenterol Hepatol. 2006;18: 545–548. doi:10.1097/00042737-200605000-00016

31. Domingo S, Solé C, Moliné T, Ferrer B, Ordi-Ros J, Cortés-Hernández J. Efficacy of thalidomide in discoid lupus erythematosus: insights into the molecular mechanisms. Dermatology. 2020;236: 467–476. doi:10.1159/000508672

32. Lu J, Liu D, Tan Y, Li R, Wang X, Deng F. Thalidomide attenuates colitis and is associated with the suppression of M1 macrophage polarization by targeting the transcription factor IRF5. Dig Dis Sci. 2021;66: 3803–3812. doi:10.1007/s10620-021-07067-2

33. Zhang F, Zhang A, Meng T, Liu X, Yang C, Liang C, et al. Epithelial redox stress programs macrophage immunometabolism through a ZNF24-MIF-NF-κB pathway in chronic nonbacterial prostatitis. Redox Biol. 2026;90: 104042. doi:10.1016/j.redox.2026.104042

34. Zhou X-L, Xu P, Chen H-H, Zhao Y, Shen J, Jiang C, et al. Thalidomide inhibits TGF-β1-induced epithelial to mesenchymal transition in alveolar epithelial cells via smad-dependent and smad-independent signaling pathways. Sci Rep. 2017;7: 14727. doi:10.1038/s41598-017-15239-2

35. Vargesson N. Thalidomide-induced teratogenesis: history and mechanisms. Birth Defects Res Part C Embryo Today Rev. 2015;105: 140–156. doi:10.1002/bdrc.21096

